# Evaluation of affinity-purification coupled to mass spectrometry approaches for capture of short linear motif-based interactions

**DOI:** 10.1101/2022.10.19.512833

**Authors:** Eszter Kassa, Sara Jamshidi, Filip Mihalič, Leandro Simonetti, Johanna Kliche, Per Jemth, Sara Bergström Lind, Ylva Ivarsson

## Abstract

Low affinity and transient protein-protein interactions, such as short linear motif (SLiM)-based interactions, require dedicated experimental tools for discovery and validation. Here, we evaluated and compared biotinylated peptide pulldown and protein interaction screen on peptide matrix (PRISMA) coupled to mass-spectrometry (MS) using a set of peptides containing interaction motifs. Eight different peptide sequences that engage in interactions with three distinct protein domains (KEAP1 Kelch, MDM2 SWIB, and TSG101 UEV) with a wide range of affinities were tested. We found that peptide pulldown can be an effective approach for SLiM validation, however, parameters such as protein abundance and competitive interactions can prevent the capture of known interactors. The use of tandem peptide repeats improved the capture and preservation of some interactions. When testing PRISMA, it failed to provide comparable results for a model peptide that successfully pulled down a known interactor using biotinylated peptide pulldown. Overall, in our hands, we find that albeit more laborious, biotin-peptide pulldown was more successful in terms of validation of known interactions. Our results highlight that the tested affinity-capture MS-based methods for validation of SLiM-based interactions from cell lysates are suboptimal, and we identified parameters for consideration for method development.

## Introduction

Protein-protein interactions (PPIs) underlie biological processes. PPIs that occur between folded globular proteins usually engage fairly large binding interfaces and these interactions are often of high affinity (1). Lower affinity interactions are commonly observed between folded globular domains and short linear motifs (SLiMs) found in the intrinsically disordered proteins or protein regions (2, 3). Here, the focus is on SLIM-based interactions. These interactions are crucial for cell function and are for example vital for the trafficking of proteins to their correct cellular localization, cell signaling, and transcriptional regulation (3, 4). SLiMs are typically 3-10 amino acid long stretches, of which a limited set of amino acids is critical for binding (4). SLiMs are, apart from their crucial role in host cellular processes, often mimicked by viral proteins to hijack host cellular functions (5). A well-known example of viral motif mimicry is the TSG101 UEV-binding PTAP motif by the GAG polyprotein of HIV, a host-virus protein-protein interaction that is critical for recruiting the ESCRT system to the location of viral budding (6). Identification of SLiM-based host-virus interactions provides detailed insights into how viruses take over the host cell and may provide insights into targets for the development of antiviral agents (7, 8).

SLiM-based interactions are challenging to capture by many high-throughput methods for interaction screening, and they are hence underrepresented in protein interaction databases such as BioGrid (9). According to estimates, the number of SLiMs in the human proteome can be up to one million, highly exceeding the number of globular domains (around 35,000) (10). The manually curated gold standard Eukaryotic Linear Motif database contains 2,222 validated human motif instances as of October 2022 (4), most of which have been found and corroborated through low-throughput experiments. However, the last decade has seen development of experimental (11-13) and computational (14, 15) approaches for proteome-wide screening of SLiM-based interactions and motif. Proteomic peptide phage display (ProP-PD) has been developed as an efficient approach for proteome-wide (and even pan-viral) screening of SLiM-based interactions (7, 16, 17). ProP-PD screening provides large-scale data on motif-based interactions from selections against a set of bait proteins. While the ProP-PD results are providing large-scale data on SLiM-based interactions, the results, like most large-scale PPI data, do require orthogonal validations. For validation of binding affinities, biophysical affinity measurements are often performed (e.g. by fluorescence polarization (FP or isothermal titration calorimetry (ITC)). Recently, higher throughput approaches such as the hold-up assay (18, 19) and spectrally encoded beads (20) have been developed, opening the door for large-scale affinity determinations of SLiM-based interactions. In addition, it is desirable that the interactions are validated by cell-based approaches such as co-immunoprecipitation (co-IP) coupled with MS-based approaches. Such analysis can be done using full-length proteins, or through pulldown with, for example, biotinylated peptides, the latter specifically validating the interactions with a SLiM-containing peptide (21, 22). One drawback with peptide-pulldown experiments is their limited throughput. Recently, PRISMA (protein interaction screen on peptide matrix) was developed as a higher throughput peptide pulldown approach (23-25). However, SLiM-based interactions may easily be lost during the washing steps of the pulldown, which may generate false negative results. Moreover, pulldown experiments coupled to mass-spectrometry will identify both binary direct interactions and indirect interactions through the pulldown of larger complexes. Thus, the results from peptide-pulldown experiments need to be treated with care in the context of identification and validation of binary SLiM-based interactions.

In this study, we systematically tested the use of biotin-peptide pulldown coupled to MS as a method for validation of SLiM-based interactions. To this end, we selected a set of known peptide ligands binding to globular domains of three different human proteins (KEAP1 Kelch, TSG101 UEV domain, and MDM2 SWIB domain) with a range of affinities (**Table 1**). We further tested how the use of single and double repeats of the SLiM-containing peptides affected the experimental outcome and compared the outcome of PRISMA in relation to classical biotin-peptide pulldown (**Figure 1**). The systematic analysis confirmed that the peptide-pulldown approaches may be used to capture some SLiM-based interactions, but also revealed important aspects to consider for capture of the known ligands.

**Table 1.**
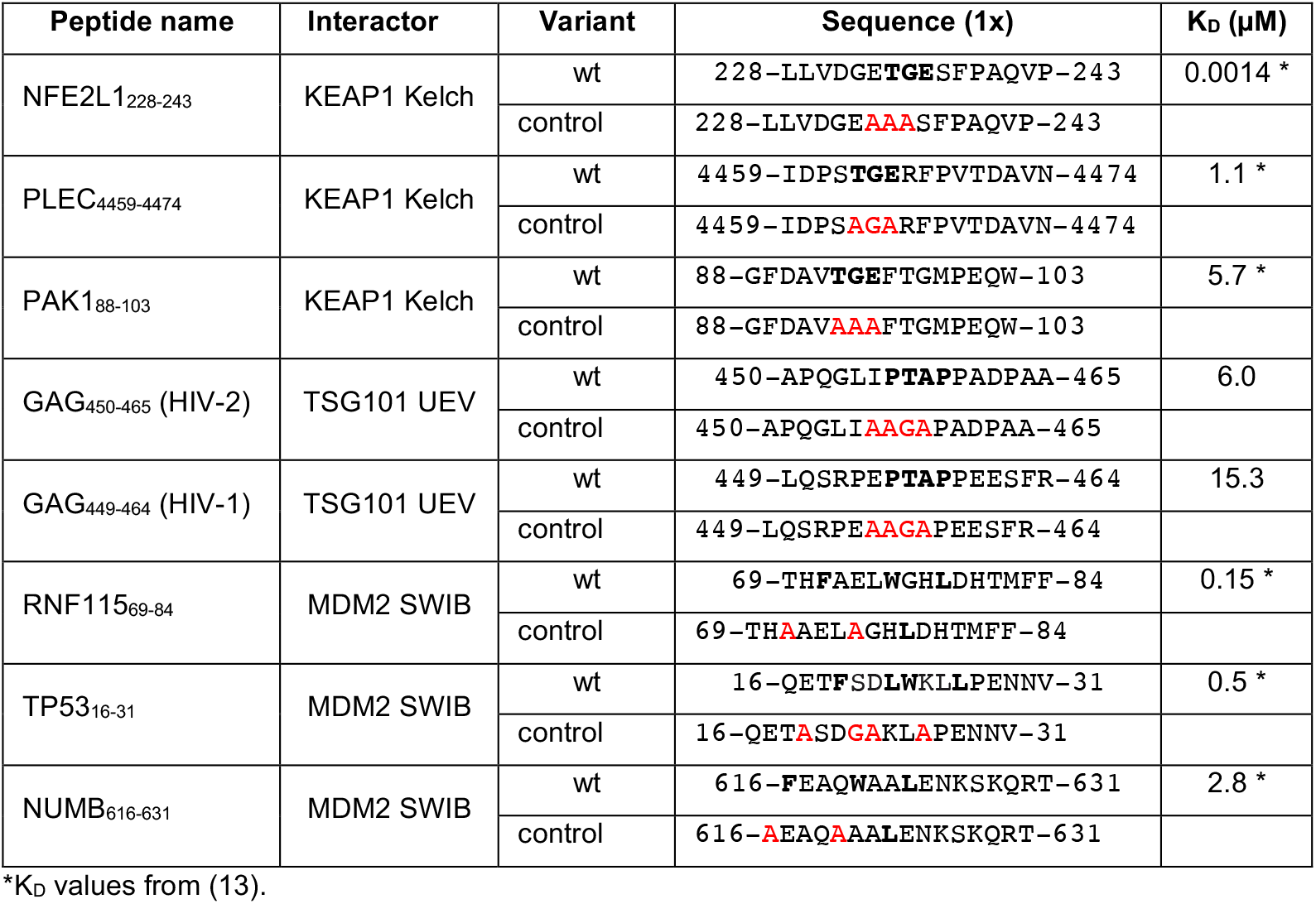
Overview of target proteins and bait peptides used. Peptide sequences (wt) are indicated (motif in bold), together with their respective negative controls (mutated amino acids indicated in red).

**Figure 1:**
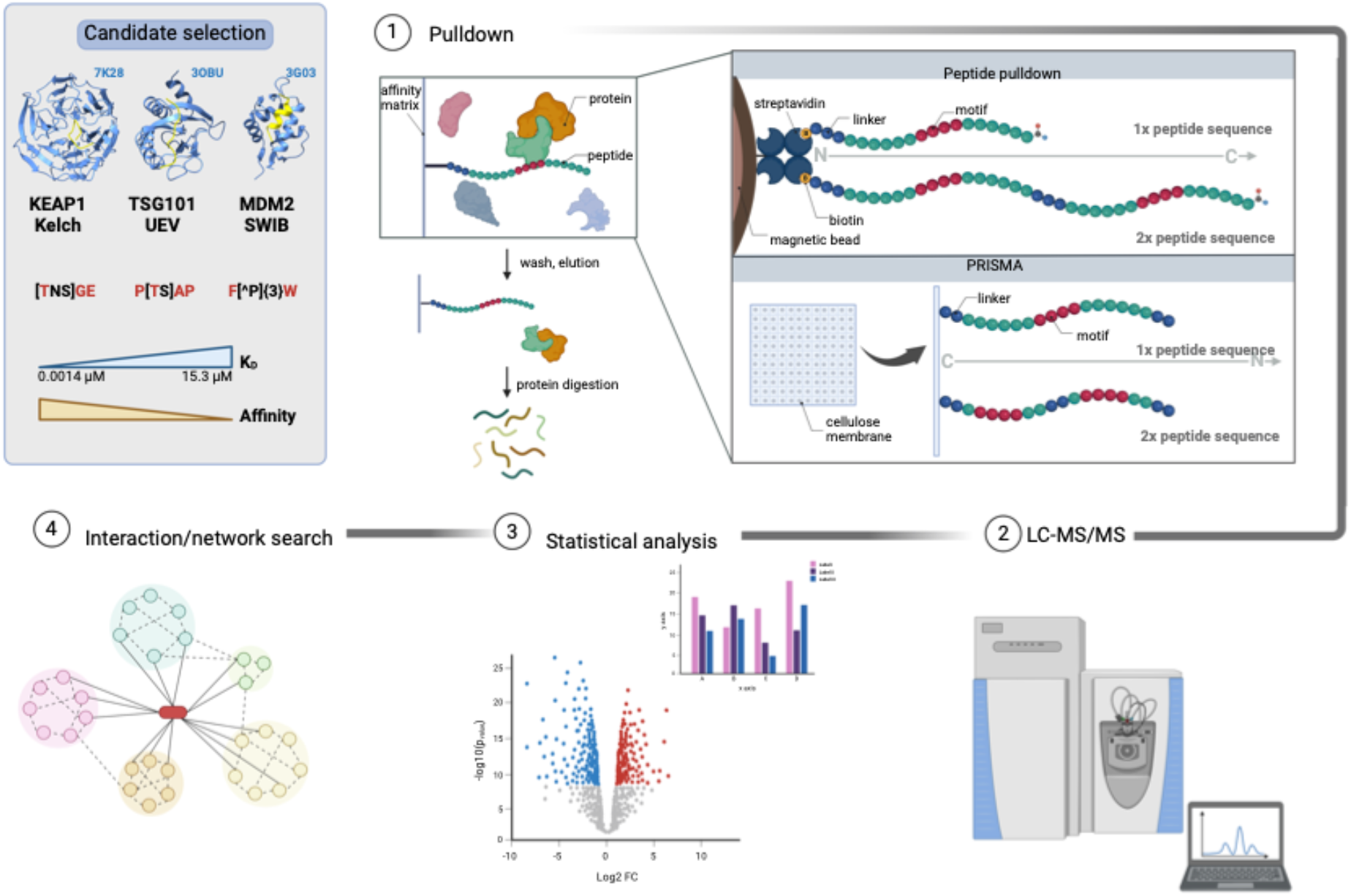
Flowchart of the peptide pulldown approaches tested, together with an overview of the data analysis pipeline. (1) Peptide ligands were selected based on previously validated interactions (13) or from ligands previously reported in ELM (4). (2) Synthetic peptides (biotinylated or spotted on a membrane) were obtained from commercial providers and used in pulldown and PRISMA experiments, where the enriched proteins are identified using LC-MS/MS. To identify meaningful interactions, statistical testing and scoring (3) were performed, together with a protein-protein interaction network search (4). (Created with BioRender.com.)

## Results

### Peptide-based affinity purification can validate previously reported SLiM-based protein-protein interactions

We aimed to explore the suitability of peptide-based pulldown to capture SLiM-based interactions. A set of motif-containing human or viral peptide sequences (19 amino acids long, including a flaking N-terminal TGS linker) that bind to human protein domains based on peptide ligands previously identified through ProP-PD selections and validated through FP-based affinity measurements (13), or based on available literature (26-28) were selected to serve as model bait peptides. The peptides were selected to include three different types of binding motifs: a short polar TGE motif binding to KEAP1 Kelch, a slightly longer PTAP motif binding to the UEV domain of TSG1, and a long helical motif binding to the SWIB domain of MDM2. Ligands were also selected to cover a range of affinities (K_D_ ranging from 0.0014 to 18.5 µM (13); **Table 1**). To explore whether having double peptide repeats could contribute to improve the outcomes of the pulldowns we also used longer peptides (38 AAs, including linkers) where the motif-containing peptides were presented in tandem. We then evaluated the results in terms of successful identification of the expected binder, its interaction partners, as well as if there were other potential interactions. In addition, for one of the KEAP1 ligands, we examined if the higher-through PRISMA method (24, 25) could be used to capture the target protein. The immobilized peptides (on beads or on a cellulose membrane) were used to pull down binding proteins from cell lysate, non-binding proteins were washed away, and the remaining proteins were digested into peptides before the LC-MS/MS analysis. Evaluation of the MS data was performed to reveal significantly enriched proteins (permutation-based FDR <0.05, S_0_:0.1) and proteins for which there was a significant SAINT score difference (>0.85) (29, 30) between the wild-type peptide and control peptide pulldown, as described on a case-by-case basis below.

### Biotin-peptide ligands consistently pulldown KEAP1

KEAP1 is a substrate adaptor of the BTB-CUL3-RBX1 E3 ubiquitin ligase complex and targets the complex to its substrates by binding to TGE-containing degradation motifs (31). We used three TGE-containing peptides from NFE2L1, PLEC, and PAK1, spanning an affinity range from nM (NFE2L1_228-243_) to low µM (PAK1_88-103_) (**Table 1**), to explore if we could use the biotin-peptide pulldown to capture KEAP1 from cell extract. We also performed the same experiments with double repeats of the peptides (sequences in **Table S1**) to explore if this would improve the capture of the target. For the PAK1 peptide, it was not possible to obtain a tandem control peptide due to synthesis problems, therefore the results from the single repeat control peptides were used for comparisons in this case. In all cases, the known interactor protein KEAP1 was observed in each replicate and in none of the control samples with mutated motifs, showing that the TGE motif is crucial for the interactions and that the biotin-peptide pulldown worked well for this target (**Table S4-6**). While the single repeat peptides enriched only a few proteins (including KEAP1), the double repeat peptides pulled down many more proteins, of which a small fraction was known ligands of KEAP1 or known interactors of the full-length proteins containing the TGE motif (NFE2L1, PLEC, or PAK1) (**Figure 2 A-C**). Looking at the overlap of proteins pulled down with the single and double repeat peptides, KEAP1 was the only common ligand. Notably, both PLEC_4459-4474_ and NFE2L1_228-243_ double repeats enriched the previously KEAP1 interactors NPM1 (32) and CSNK2A1 (33). The PAK1 peptide (in single and double repeats) pulled down a set of other proteins (RPA1/2/3 and BANF1) with high confidence, and they may thus represent novel interactors of PAK1.

**Figure 2:**
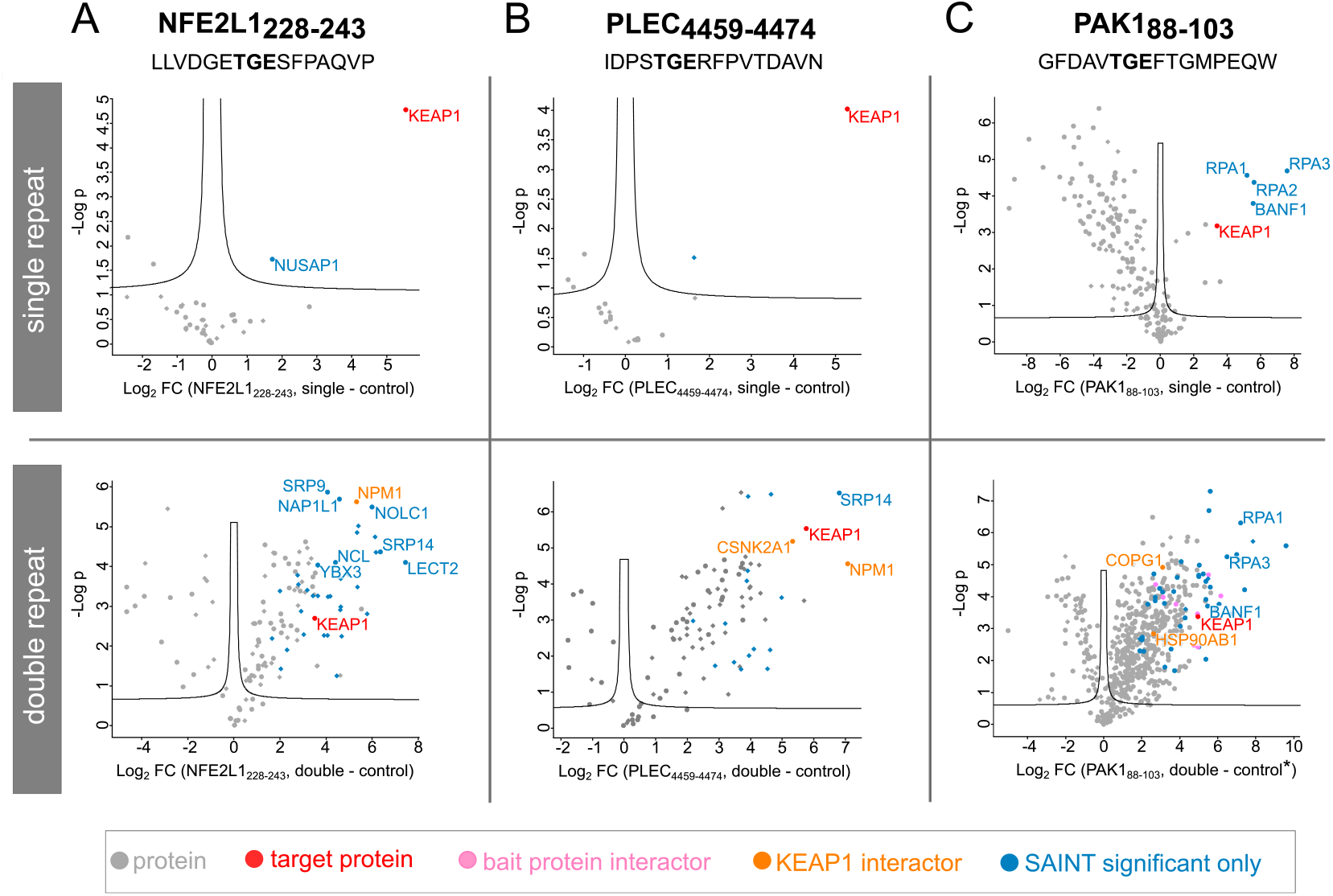
Volcano plots of peptide pulldowns using TGE motif-containing peptides. (permutation-based FDR<0.05 (250 permutations), S_0_:0.1, SAINT significant: SAINT score >0.85.) Top panel: single peptide repeat, bottom panel: double peptide repeat. A) NFE2L1 peptide, residues 228-243. B) PLEC peptide, 4459-4474. C) PAK1 peptide, residues 88-103.

Taken together, the KEAP1 results showed that the TGE-containing peptides efficiently pulled down the expected target. The single repeat peptide pulldown produced the cleanest data on binary interactions, while the double repeat also facilitated the enrichment of known interactors of the prey protein, as well as many other proteins.

### A tandem PTAP motif is necessary for the capture of TSG101

We next explored if we could use PTAP motif-containing peptide pulldown to capture TSG101. In this case, we used two TSG101 UEV binding peptides from GAG (Human immunodeficiency virus type 1 (HIV-1)) and GAG (HIV-2) as model peptides (27, 34). For these two peptides, we first determined the K_D_ values for their interactions with the TSG101 UEV domain (**Figure 3A-B**, K_D_ 15.3 and 6.0 µM for GAG (HIV-1) and (HIV-2), respectively). Single and double repeats of the peptides were used in peptide-pulldown experiments. LC-MS analysis revealed a low number of enriched proteins, and neither of the single repeat peptides pulled down TSG101 (**Figure 3C-D; Table S4-6**)). In contrast, the double repeat GAG (HIV-1) peptide (but not GAG (HIV-2)) successfully pulled down TSG101, together with several other members of the ESCRT-1 complex (MVB12A, VPS28, VPS37B, VPS37C) (6, 35). The linkage of two PTAP motifs thus improved the capture of TSG101, which can be explained by the increased local concentration of the SLiM. The interaction between the PTAP motif and TSG101 appears to be very specific, based on the LC-MS analysis. This might be connected to the fact that the TSG101 UEV domain has a marginally different structure than the few other UEV-domains present in human proteins (36).

**Figure 3:**
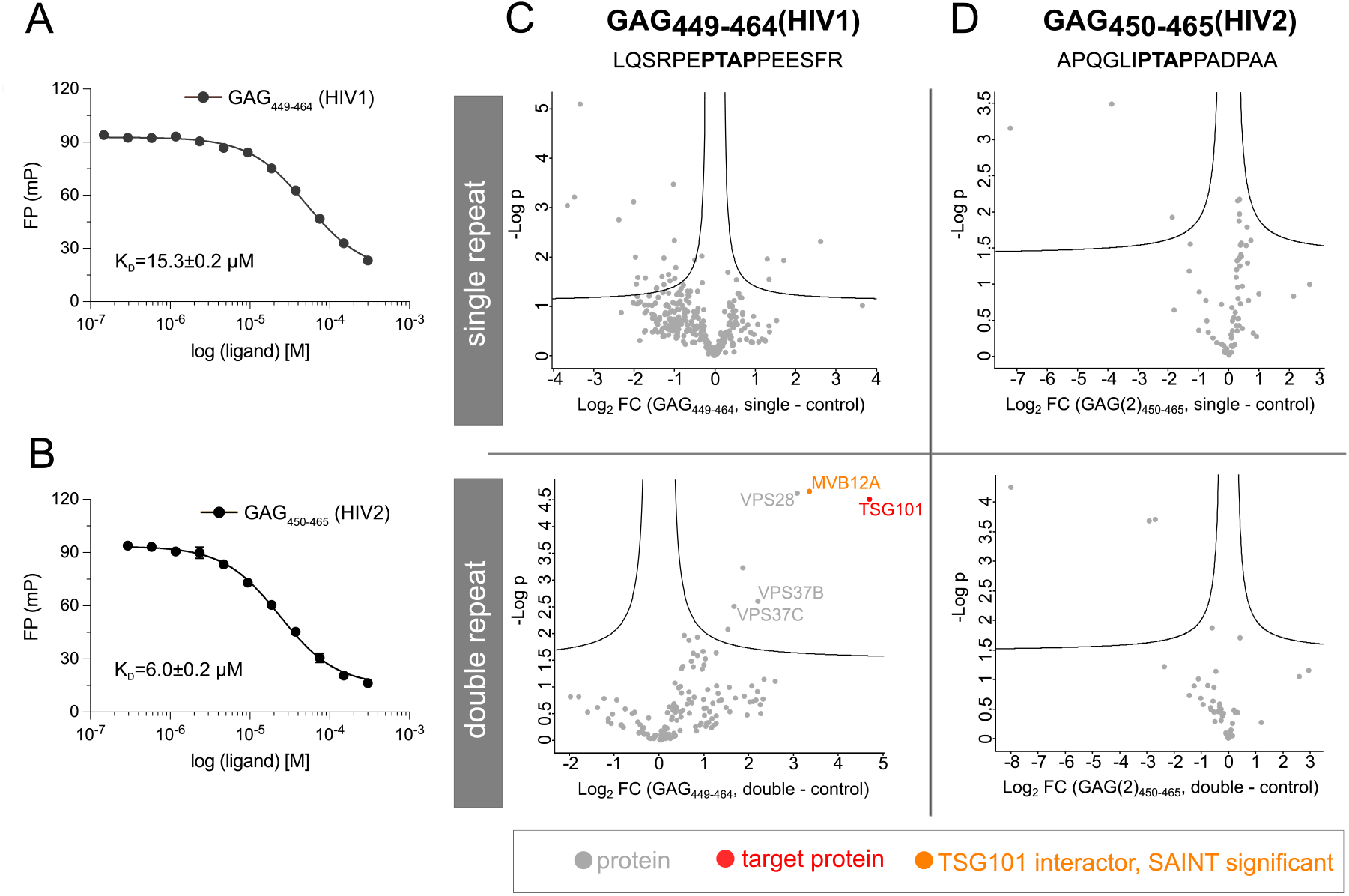
A tandem PTAP motif is necessary for biotin-peptide pulldown of TSG101. A-B) Fluorescence polarization monitored displacement experiments of TSG101 UEV domain binding to A) GAG (HIV-1) residues 449-464 and B) GAG (HIV-2) peptide, residues 450-465, (indicated with error bars showing one standard deviation). K_D_ indicated with one SEM (standard error of the mean). C-D) Volcano plots of peptide pulldowns using PTAP motif-containing peptides (permutation-based FDR<0.05 (250 permutations), S_0_:0.1, SAINT significant: SAINT score >0.85). Top panel: single peptide repeat, bottom panel: double peptide repeat. C) GAG (HIV-1) peptide, residues 449-464. D) GAG (HIV-2) peptide, residues 450-465.

Taken together, we find that biotin-peptide pulldowns paired with LC-MS analysis can be used to identify and validate SLiM-based interactions and that the robustness of the approach can be improved by presenting the motifs in tandem, as shown for the TSG101 binding peptides.

### MDM2 binding peptides capture a large number of other proteins

The E3 ubiquitin-protein ligase MDM2 has a SWIB domain that binds to a helical degradation motif which folds into an alpha-helical structure upon binding (37). Among its well-known ligands is the binding to the cellular tumor antigen p53 (p53), an interaction that leads to proteasomal targeting of p53 and inhibition of p53-mediated cell cycle arrest (37). We used MDM2 SWIB binding peptides from RNF115, p53, and NUMB (K_D_ values ranging from 0.15-2.8 µM (13)) as baits in peptide pulldowns against HaCat cell lysates, using single and double repeats of the peptides as before. From these pulldowns, a large number of proteins were found to be pulled down (**Figure 4, Figure S1, Table S2**). However, the expected binder MDM2 was not identified in any of the samples. The affinities of the interactions between the bait peptides and MDM2 SWIB are similar to the affinities of the KEAP1 Kelch peptide interactions tested (**Table 1**), suggesting that the failure to pulldown MDM2 may be explained by other factors, such as a low concentration of MDM2 in unstressed cells (37) or competition for the binding motif by other binding partners. Indeed, the p53 MDM2 binding motif overlaps with the p53 transactivation domain, which can be bound by the TAZ1 and TAZ2 domains of p300 and CBP (38), as well as by Bcl-2 and Bcl-X(L) (39). However, none of these proteins were significantly enriched in the pulldown although a large number of other proteins were identified (**Figure 4A; Table S4-6**). In the cases of RNF115_69-84_, a large number of significantly enriched and SAINT-significant proteins were pulled down, with ATN1 as the most prevalent known interactor of RNF115 potentially suggesting a direct interaction between the RNF115 peptide and ATN1. The NUMB_616-631_ peptide proved to be less promiscuous, resulting in only a few SAINT significant proteins.

**Figure 4:**
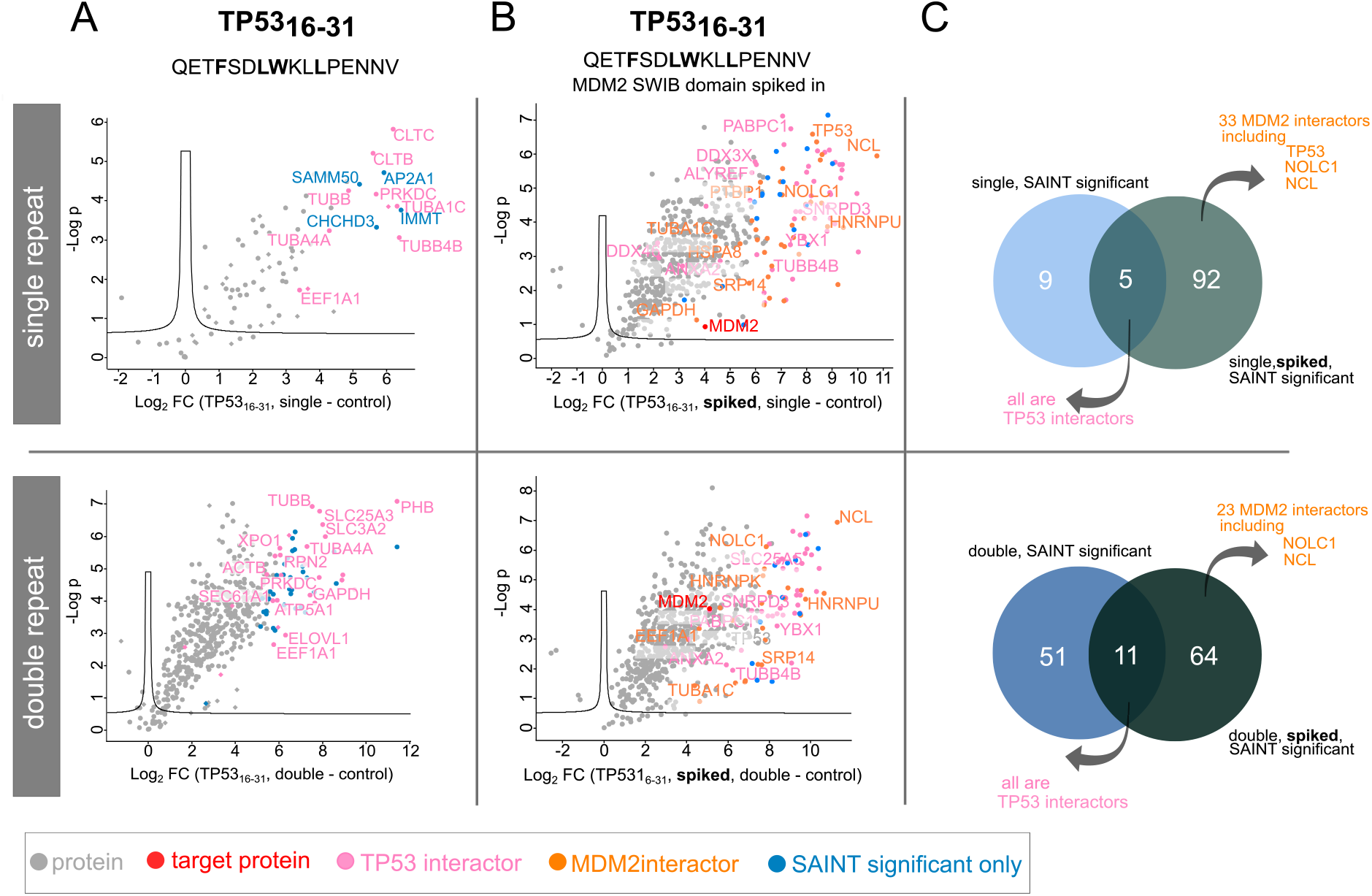
Volcano plots of peptide pulldowns using p53 peptides. Top panel: single peptide repeat, bottom panel: double peptide repeat. A) Biotin-peptide pulldown using p53 peptide (residues 16-31) and negative control. B) Same as in A) but with MDM2 SWIB domain spiked in the cell lysate. C) Venn diagram of SAINT significant proteins without or with MDM2 SWIB domain spiked into the cell lysate. (Permutation-based FDR<0.05 (250 permutations), S_0_:0.1, SAINT significant: SAINT score >0.85.)

To test if the failure to detect MDM2 in the biotin-peptide pulldown was due to a low MDM2 concentration we spiked in recombinantly expressed GST-tagged MDM2 SWIB domain in the cell lysate and carried out the pulldown with p53_16-31_. In this case, MDM2 was identified and quantified in every sample, including the controls, together with several known p53 interactors, as well as a large number of other proteins. Notably, however, MDM2 was significantly enriched in both the single and double repeat samples compared to their respective controls, by 3.8 and 33.4 times higher average LFQ intensities, respectively. Thus, one explanation for the failure to detect MDM2 in the pulldowns may be a low cellular MDM2 concentration

### Evaluation of PRISMA for capturing SLiM-based interactions

The analysis of the KEAP1 Kelch and TSG101 UEV1 domain binding peptides confirmed that these proteins can be pulled down from cell lysate using biotinylated peptides. A drawback in the pulldown approach of using biotinylated peptides captured with streptavidin-coated beads is the relatively high cost of the synthetic peptides and the time-consuming sample preparation. During the last couple of years, PRISMA has been developed as a more economical, and less laborious approach (23, 24). We therefore decided to evaluate the biotin-peptide pulldown in comparison to the PRISMA. We selected the KEAP1 Kelch binding PLEC_4459-4474_ peptide as a model peptide as it robustly pulled down KEAP1 in the biotin-pulldown both as single and tandem repeats. Following the same approach as above, we design a PRISMA array with single and double repeats of the TGE-containing peptide and used motif-mutated peptides as negative controls. Due to the peptide length restriction set by the PRISMA array (20 amino acids), the double repeat peptide was trimmed down to 7 amino acid-long sequence covering the core motif, which was then linked in tandem (20 amino acids, with two repeats and linkers). This limits the range of SLiMs that can be investigated as double repeats, as some motifs are more dispersed.

We tested two versions of a previously described protocol (25) for the PRISMA array analysis, in the first case using stable isotope labeling by amino acids in cell culture (SILAC) quantification (40) and in the second case using unlabeled cell lysate and LFQ setup (41, 42). When following the SILAC labeling approach, we failed to identify KEAP1 as a binder of the PLEC peptides using either single or double repeats (**Table S3**). When the experiment was performed with unlabeled cell lysate and LFQ setup KEAP1 was not identified among the protein enriched for the single motif repeat peptide, but it was found as a unique hit in the PLEC_4460-4466_ double repeat samples although with low intensity (**Figure 5, Table S4**). Notably, KEAP1 would not have been identified as a relevant binder from these experiments without the a priori knowledge of the interaction. The LFQ intensities acquired by peptide pulldown and PRISMA cannot be directly compared, but it is worth mentioning that the identified sequence coverage for KEAP1 in peptide pulldown was on average 65.9% (27 identified peptides), compared to 5.8% and (3 peptides identified) in the case of PRISMA. This indicates a much lower concentration of enriched KEAP1 in PRISMA experiments compared to pulldown experiments.

**Figure 5:**
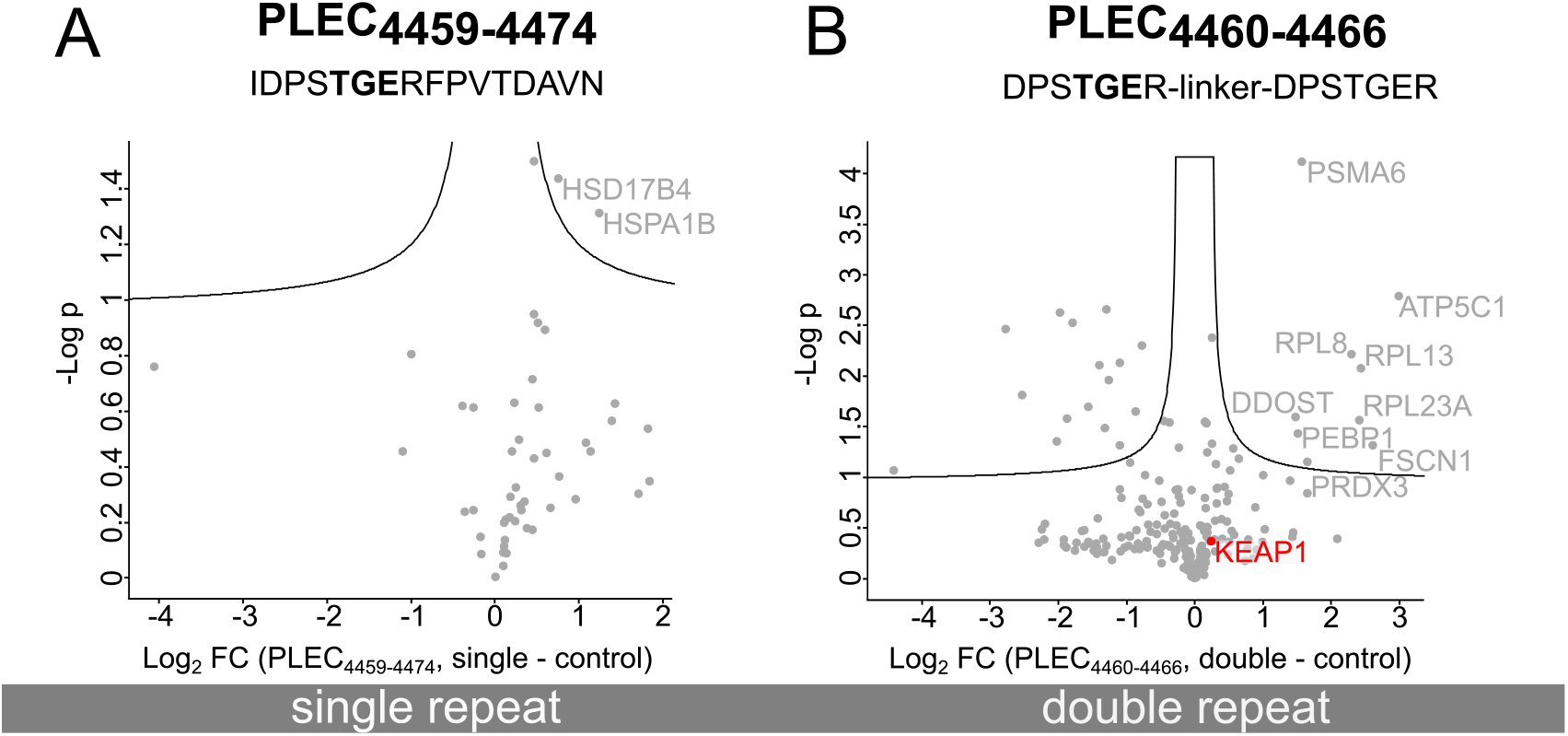
Volcano plots of label-free quantification of PRISMA assays using TGE motif-containing peptides (permutation-based FDR<0.05 (250 permutations), S0:0.1, SAINT significant: SAINT score >0.85.) Left panel: single peptide repeat (PLEC_4459-4474_), right panel: double peptide repeat.

Our analysis suggests that the PRISMA array is less reliable in terms of the pulldown of true positive SLiM-binding proteins. To ensure that there was no obvious handling mistake with the PRISMA array, we obtained an array displaying the same GLUT1 and SOS1 peptides as used in a previous study (25). Despite using another cell line, a reasonable overlap was observed between our experiments and published data supporting the validity of the outcome of our results (**Table S7**).

## 4. Discussion

We here tested peptide pulldown coupled to MS as a method for validation of SLiM-based interactions of a select set of peptides known to bind to different bait proteins. MS is especially useful in this area, as it is highly sensitive and provides unbiased identification of a high number of proteins. In addition to different ligands and target proteins, we tested how the use of single and double repeats of the SLiM-containing peptides affected the experimental outcome. The systematic analysis revealed that the single repeat biotinylated peptide pulldown may be used to capture SLiM-based interactions, as shown for KEAP1. For the TSG101 binding peptide, we found that a tandem PTAP containing peptide repeat was required to successfully pulldown the protein. Overall, the experimental results showed a higher number of proteins pulled down using the double repeat peptides. However, these pulldowns were also less specific, while the single repeat peptide provided clearer information about direct interactors, when successful (i.e. KEAP1 ligands). On the other hand, the increased local concentration arising from presenting multiple repeats of the same motif might prove to be essential to capture interactors, as clearly displayed using the PTAP motif against TSG101 UEV domain. However, we found that the approach failed to return known ligands for several cases (e.g. for MDM2), even when the binary interaction between the peptide and the target protein is known to be of higher affinity. In part, this may be explained by low target protein concentration, or by competing interactions with other proteins.

We further evaluated the applicability of PRISMA in relation to the biotinylated peptide pulldown using a validated KEAP1 Kelch domain binding peptide as a model ligand. The PRISMA approach has been proposed as an elegant alternative to the biotin-peptide pulldown (23-25), with the peptides spotted on the membrane offering an economical alternative in comparison to the biotinylated peptides, and a less laborious protocol for upstream processing. However, in our hands and with the peptide tested, we found it challenging to capture the known interactors which had been successfully identified using peptide pulldown. This might be the result of lower purity peptides synthesized on the membrane, as well as the more complex digestion environment. Making the choice between the two methods, and with a limited number of ligands to be tested, we would favor the biotinylated peptide pulldown. Several factors contribute to making it challenging to capture SLiM-based interactions using peptide pulldowns, including rapid dissociation of this kind of interaction in combination with the multiple washing steps during the pulldown. Other limitations are related to what peptides can be synthesized (length and composition). For PRISMA the length restriction sets a limit of 20 amino acids. Longer peptides can be used if using biotinylated peptides instead, but the synthesis of longer peptides may be challenging, and longer peptides may also become fairly costly to obtain. One way to circumvent the need to obtain synthetic peptides is to genetically fuse them to a tag (eg. GFP) and express them intracellularly (7, 17). A drawback with these approaches is that the presence of the peptides in the cell may inhibit the normal cellular function of the binding protein, which, in some cases, may perturb cell survival and cell growth. The size, type, and location of the used tag can also influence function and interaction.

In summary, we conclude that the pulldown-based validations of SLiM-based interactions remain challenging and that other approaches for higher throughput validations are desirable, especially given the rapidly growing number of predicted interactions.

## Experimental procedures

### Cell culture and lysis

HaCat cells were cultured in DMEM+GlutaMAX + 10% FBS (Gibco). Cells close to confluency were washed with PBS and lysed with lysis buffer (1% NP-40 substitute (Sigma 74385) in PBS + protease inhibitors). The cells were scraped and agitated in a lysis buffer for 20 minutes on ice. The lysate was centrifuged at 16000 *g* for 10 minutes at 4°C. The supernatant was collected and stored at −80°C. To achieve a high protein concentration for PRISMA smaller lysis buffer volumes were used.

SILAC-labeled HaCat cells were generated using DMEM for SILAC (Thermo, 88364) +10% FBS (Gibco, A3382001). For the heavy labeled cells, 0.89 M Lys8 (Silantes, 211604102) and 0.40 M Arg19 (Silantes, 201604102) were added, while L-lysine (Sigma, L8662) and L-Arginine (Sigma, A6969) were used for the light labeled cells. The cells were kept in these media for at least 6 cell doublings, and no trypsin was used during cell passaging. The cell lysates were prepared as shown above. Separate aliquots of cells were prepared for label incorporation check, according to a detailed practical guide (43). The incorporation rate of heavy amino acids was verified using a freely available R script (https://github.com/BerndHessling/MaxQuant-Incorporation-Control).

### Peptide pulldown

Peptides (>90% purity; **Table S1**) used in peptide pulldown experiments were N-terminally biotinylated and C-terminally amidated (GeneCust). The peptides were dissolved to ca. 1 mg/ml concentration in phosphate buffered saline (PBS) (Sigma 4417, 0.01 M phosphate buffer, 0.0027 M potassium chloride, and 0.137 M sodium chloride, pH 7.4). The pH was adjusted with ammonium bicarbonate solution as needed to dissolve the peptides. Peptide aliquots were stored at −80°C. The cell lysate was diluted to 0.8 mg/ml total protein concentration and 0.47% NP40 using dilution buffer (PBS plus protease inhibitor cocktail (Roche 04693159001) and precleared using Invitrogen™ Dynabeads™ MyOne™ Streptavidin T1 magnetic beads for 2-4 hours at 4°C during gentle rotation. Peptide stock solutions were diluted with washing buffer (0.1% NP40 substitute (Sigma 74385) in PBS) to 0.025 mg/ml (single repeat) or 0.05 mg/ml (double repeat). Peptides (500 µl of 0.025 mg/ml stock solution) were immobilized on the surface of 75 µl streptavidin-coated magnetic beads by incubating for 1 hour at room temperature while shaking. Non-bound peptides were removed by washing three times with 1000 µl washing buffer and then 750-912 µl pre-cleared lysate was added to the beads. The pulldown proceeded at 4°C overnight while the samples were rotated. The samples were then washed five times with ice-cold PBS and the bound proteins were eluted by incubation for 5 minutes with 200 µl 0.1 M glycine-HCl, pH 2.5 at room temperature while rotating. The eluate was neutralized in a fresh tube using 25 µl 1 M ammonium bicarbonate. Samples were flash-frozen in liquid nitrogen and stored at −80°C until further processing. After thawing, the samples were reduced using 1 µl 1 M DTT and incubated at 50°C for 15 minutes while shaking, alkylated using 1 µl 550 mM iodoacetamide (IAA), and incubated in darkness for 15 minutes at room temperature. 88 µl 50 mM ammonium bicarbonate was added to dilute the sample and 2 µl of 0.2 µg/µl trypsin was added to digest the proteins at 37°C overnight while shaking. To stop the digestion the solution was acidified to a pH<3 using a small volume of 83.3% AcN, 16.7% TFA. In-house-made STageTips (44, 45) were used to remove salts. In brief, 2 layers of a C18 membrane (3M Empore) were packed into 200 µl pipette tips. The membranes were wetted using methanol, washed with 80% AcN, 0.1% formic acid then washed twice with 0.1% formic acid in water. The acidified samples were loaded onto the membrane and washed with 0.1% formic acid in water. Finally, the captured peptides were eluted using 80% AcN, 0.1% formic acid, then vacuum-dried and stored at −80°C. For the spike in samples, GST-tagged MDM2 SWIB domain was added to the cell lysate to a final concentration of 4 µM (ca. ten times the K_D_ (207.8 µl SWIB domain (7.41 mg/mL in 50 mM Tris, 150 mM glutathione pH 8.0) to make 11 ml spiked lysate)). For the peptides, GAG_449-464_ (HIV-1) and GAG_450-465_ (HIV-2) the protocol contained the following differences: the cell lysate was diluted to 0.8 mg/ml protein concentration with 0.29% NP40 concentration. 0.025 mg/ml peptide concentration was used in the case of both single and double repeat peptides. Three 10-minute-long washing steps were used to remove unbound peptides after peptide immobilization. 1250 µl diluted lysate was used per sample. Before elution, 3-5 washing steps were carried out with 1000 µl BW1.

### PRISMA

PRISMA membranes were acquired from JPT Peptide Technologies GmbH with the N-terminal ends of the peptides acetylated (**Table S3**). The method used for the pulldown is described in detail elsewhere (25). Briefly, the PRISMA membranes were incubated with PBS for 15 minutes at room temperature while shaking, followed by incubation with cell lysate in plastic pouches for 20 minutes, 4°C while rotating. Membranes were washed three times for 5 minutes in 25-30 ml ice-cold PBS (on ice). Membranes were placed on a glass tile and dried at room temperature. The spots were punched out using a 2 mm biopsy punch and placed in a well of a 96-well plate. For the SILAC experiments, the corresponding sample and control were placed in the same well. In the following, amounts show what was used for LFQ experiments (one spot per well)/SILAC experiments (2 spots per well). 20/40 µl denaturation buffer (6M urea, 2M thiourea, 10mM HEPES) was added to each well. The proteins were reduced with 2/4 µl of 10 mM DTT in each well for 30 minutes at 37°C and alkylated with 2/4 µl of 55 mM IAA for 45 minutes at room temperature in darkness. The samples were diluted with 100/150 µl 50 mM ammonium bicarbonate and the proteins were digested with 0.5/1 µg trypsin and 0.5/1 mAU LysC overnight at 37°C. Samples were acidified by adding 4/6 µl of 25% TFA. From here, the samples were processed as in the case of the peptide pulldowns.

### LC-MS/MS

The LC-MS/MS system consisted of an Easy-nLC 1000 nano-HPLC (Thermo Fischer Scientific), and Q Exactive Plus mass spectrometer (Thermo Fischer Scientific) with an EASY-Spray electrospray (ESI) ion source. The liquid chromatography system contained an Acclaim PepMap 100 pre-column (Thermo Fischer Scientific, 75 µm x 2 cm, 3 µm, 100Å) and an EASYspray PepMap RSLC C18 analytical column Thermo Fischer Scientific, EASYspray, 75 µm x 15 cm, 2 µm, 100Å).

A gradient LC method going from 4-76% acetonitrile in 79 minutes was used for peptide separation. The mass spectrometer was used in the positive ion mode with the ESI spray voltage of 1.9 kV, full scan resolution was set to 140000 (400-1700 m/z) and the MS/MS resolution was 17500 while the used automatic gain control (ACG) target was 3×10^6^ and 1×10^5^, for MS and MS/MS respectively. A data-dependent approach was used, where the top 10 most abundant ions were fragmented and further analyzed, also employing a dynamic exclusion of 30 s.

The dried peptides were resuspended in 21 µl 0.1% formic acid and 5 µl of this solution was analyzed in each case.The SILAC samples were then diluted 2 times with 0.1% formic acid, and 5 µl of this solution was analyzed.In the case of the peptides, GAG_449-464_ and GAG(2)_450-465_ an Acclaim PepMap RSLC column was used (Thermo Fischer Scientific 164940, 75 µm x 15 cm, nanoViper, C18, 2µm, 100Å) for separation complete with a Nanospray Flex ion source. The gradient went from 4 to 80% acetonitrile in 79 minutes. Controls were processed the same way as pulldowns with wt peptides.

### Data analysis

The data were searched in MaxQuant (2.0.1.0) against the FASTA file of the human proteome (Uniprot, UP000005640 *Homo sapiens* reference proteome, downloaded 2021.04.30., 20626 entries). For this search, both the single and double repeat baits and their corresponding controls were analyzed together. Label-free quantification was selected (MaxLFQ algorithm (41)) with a minimum ratio of 2. Minimum 2 identified peptides and minimum one unique peptide were required for protein identification. For fixed modification carbamidomethylation of cysteines was selected, while methionine oxidation and N-terminal acetylation were allowed as variable modifications. Trypsin/P was used as the digestion enzyme with maximum 2 missed cleavages allowed. PSM and protein FDR were set to 0.01. The first and main search peptide mass tolerances were 20 and 4.5 ppm, respectively. In the case of SILAC experiments, the multiplicity was set to 2 and heavy labels Arg10 and Lys8 were selected. The re-quantify option was enabled. The other settings were the same as above.

Significantly enriched proteins were determined using Perseus (2.0.3.0) (46) using the MaxQuant output file and selecting the LFQ intensity values as main. Potential contaminants, proteins only identified by site or reverse hits were removed. The LFQ intensity values were transformed using a log2(x) base. The proteins were further filtered, only keeping those that were quantified in at least 3 out of the 3 replicates (own PRISMA: 2 out of 2) in at least one group (sample and control). The missing values were replaced by imputation, using a normal distribution with a width of 0.3 and down shift of 1.8, selecting the total matrix option. Following filtering, a two-sided t-test was carried out with a permutation-based FDR approach (250 permutations, FDR <0.05, S0:0.1), and the data were visualized in volcano plots.

The same evaluation was done with PRISMA results from (25) (**Table S7**), using imputed LFQ data as main, removing empty rows and carrying out the t-test.

Known interactors of the proteins of interest were downloaded from BioGRID 4.4 (9) (accessed 2022.06.01.) and filtered for *Homo sapiens* and also HIV in the case of the GAG/TSG101.

Significance analysis of interactome (SAINT) (29, 30) scoring was carried out for each dataset. This tool requires input about baits, controls, and LFQ intensity information and generates a confidence score. The input files were generated using an in-house written script and a precompiled library of SAINTexpress v3.6.3. from the SourceForge website (https://sourceforge.net/projects/saint-apms/files/) was employed using the intensity option. Interactors with SAINT scores higher than 0.85 were considered significant.

### Protein expression and purification for fluorescence polarization

pHH0103 vector encoding human TSG101 UEV (1-145) domain was described previously (13). The protein for affinity measurements was expressed as N-terminally tagged 6xHis-GST-Thrombin cleavage site, fusion constructs in E. coli BL21(DE3), using 2YT media (16 mg/mL peptone, 10 mg/mL yeast extract, 5 mg/mL NaCl) including selective antibiotics. The incubation at 37°C continued until an OD600 of 0.6 was reached. Protein expression was then induced with 1 mM IPTG (isopropyl β-D-1-thiogalactopyranoside). For protein expression, the incubation was continued at 30°C for 4 hours or at 18°C overnight. After harvesting the bacterial cultures (centrifugation at 4,500 *g* for 10 minutes and 4°C), the pellet was resuspended in lysis buffer (50 mM Tris/HCl pH 7.8, 300 mM NaCl, 10 µg/mL DNase I and RNaseA, 4 mM MgCl_2_, 2 mM CaCl2 and cOmplete EDTA-free Protease Inhibitor Cocktail) and the cells were lysed using a cell disruptor at 1.7kBar. After separating the pellet (centrifugation at 20,000 *g* for 40 minutes) the supernatant was sterile filtered using a 0.2 µm PES filter. The filtered supernatant was applied to Glutathione Agarose (Pierce), and purified according to the manufacturer’s instructions. The eluate was further processed by enzymatic cleavage of the 6xHis-GST tag using thrombin overnight at 4°C. The resulting sample was incubated with nickel Sepharose excel resin and the protein of interest was acquired from the unbound phase. Buffer exchange was performed using a HiPrep 26/10 desalting column to 50 mM potassium phosphate buffer pH 7.5. The protein was subjected to SDS-PAGE gel electrophoresis and matrix-assisted laser desorption/ionization time-of-flight mass spectrometry (MALDI-TOF/MS) to confirm identity and purity. The concentration was measured based on absorbance at 280 nm.

### Fluorescence polarization

FITC-labeled and unlabeled peptides were ordered from GeneCust (>95% purity). The fluorescence polarization experiments were carried out in 50 mM potassium phosphate buffer pH 7.5. Black, flat bottom 96-well non-binding surface plates were used and the measurements were carried out at room temperature using a SpectraMax iD5 plate reader and the excitation/emission wavelengths 485/535 nm. The G-factor was adjusted using wells only containing the FITC-labeled peptide so that the background fluorescence polarization value would be between 10-40 mP. The affinity between the FITC-labeled peptides and the purified TSG101 has been determined previously using saturation binding experiments (17). To determine the affinities of the unlabeled peptides the displacement experiments were performed, where the constant concentration of FITC-labeled peptide (10 nM) and TSG101 UEV (8 μM) was challenged with increasing concentration of unlabeled displacer peptide (1:1 dilution series with the highest concentration of the peptides being 300 μM). From displacement experiments, IC50 values were obtained using a sigmoidal dose-response fitting model (allowing for variable slope; GraphPad Prism 9, version 9.4.1). The IC50 values were further converted to *K*_*D*_ values for the displacing peptides as described previously (13). All experiments were performed in at least 3 technical replicates.

### Protein expression and purification (MDM2 SWIB domain for spiking in)

The human MDM2 SWIB domain was expressed and purified as previously described (13). Briefly, the domain was expressed as a GST-tagged construct (pETM33 plasmid) in BL21 Gold (DE) E. coli bacteria. Bacteria were grown in 2YT media and expression was induced at 0.7-0.8 OD_600_ with 1 mM isopropyl b-D-1-thiogalactopyranoside. After that, the bacteria were pelleted (3,500 *g* for 15 min) and pellets stored at −20 °C. The pellet was resuspended in lysis buffer (0.05% Triton-100, 5 mM MgCl_2_, 10 µg/mL DNase in phosphate buffered saline (PBS), complemented with cOMplete EDTA free protease inhibitor and lysozyme), incubated for 1 h in the cold room while shaking and sonicated (Sonics Vibracell) in the meantime (2 s pulse, 2 s pause, 20 seconds total, 70%). The suspension was then centrifuged for 1 h at 16,000 *g* at 4 °C and the supernatant was added to 1 mL of glutathione beads (GE Healthcare) equilibrated in PBS. After incubation with the beads for 1 h in the cold room while shaking, the beads were washed 5 times with 15 mL PBS by spinning the beads down after each wash for 5 min at 500 *g*. Protein was eluted in 1 mL fractions in 50 mM Tris pH 8.0, 10 mM glutathione. The concentration was determined by measuring the absorption at 280 nm with a NanoDrop (ND-1000).

## 5. Data availability

The mass spectrometry proteomics data have been deposited to the ProteomeXchange Consortium via the PRIDE (47) partner repository with the dataset identifier PXD037380.

## 6. Acknowledgement

This work was supported by the grants from the Swedish Foundation for Strategic research (Y.I., P.J., S.B.L.: SB16-0039). We thank Evangelia Petsalaki for her input on data analysis.

## 8. Author contributions

E.K: Conceptualization, Methodology, Formal analysis, Investigation – Cell culture, peptide pulldowns, PRISMA and LC-MS/MS, Visualization, Writing-Original draft preparation S.J.: Investigation – Protocol development. Peptide pulldown. F. M.: Investigation – Fluorescence polarization and protein expression, Writing - Review & Editing. L.S.: Software. J.K.: Investigation – Protein expression and purification. **P. J**.: Supervision, Resources, Funding acquisition. S. B. L.: Supervision, Conceptualization, Resources, Funding acquisition, Writing - Review & Editing. Y. I.: Supervision, Conceptualization, Resources, Funding acquisition, Writing - Review & Editing.

## Supplemental information

### Supplemental figure

**Figure S1:**
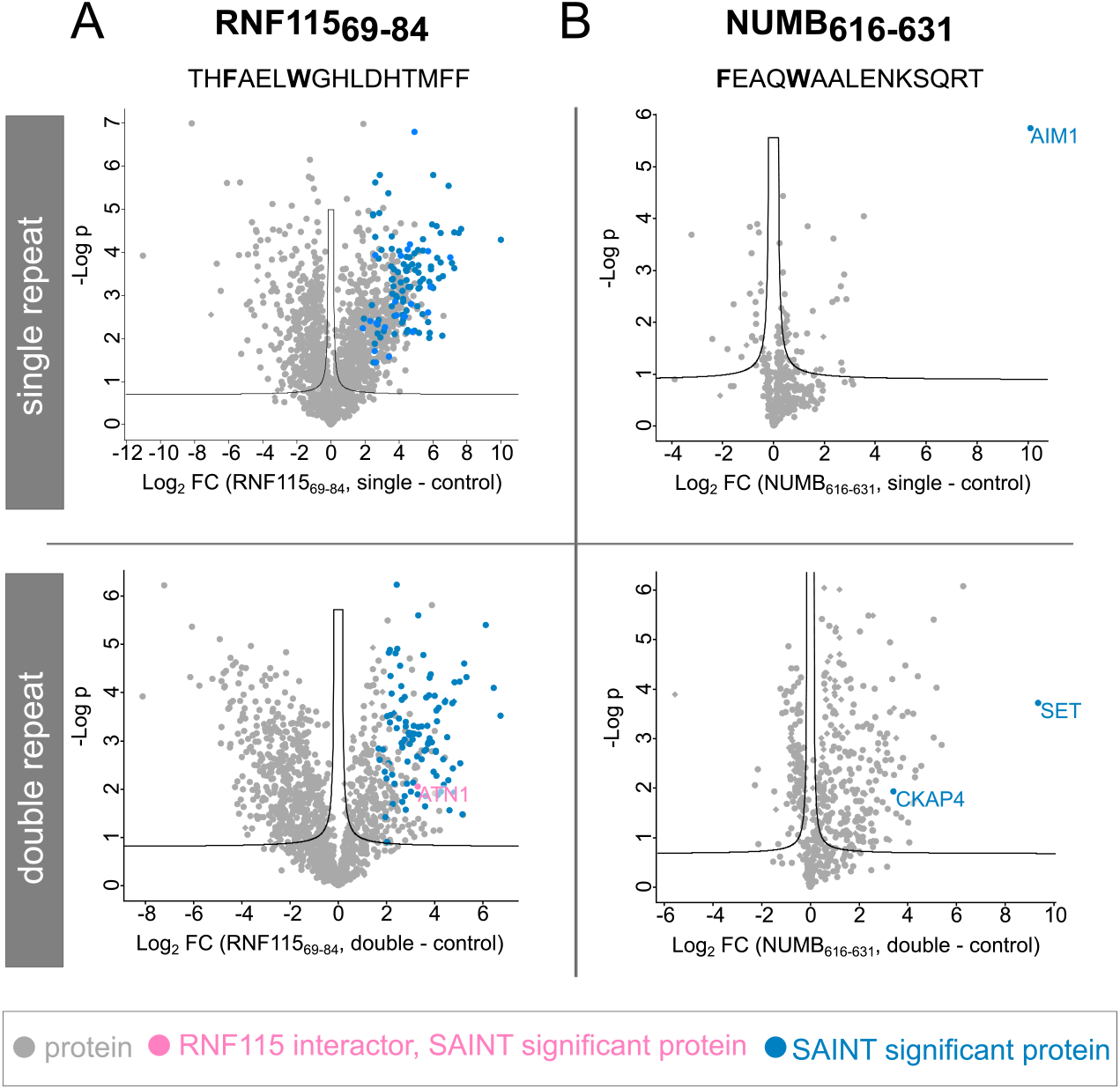
Volcano plots of peptide pulldowns using MDM2 binding peptides from RNF115 and NUMB (permutation-based FDR<0.05 (250 permutations), S_0_:0.1, SAINT significant: SAINT score >0.85.) Top panel: single peptide repeat, bottom panel: double peptide repeat. A) NFE2L1 peptide, residues 228-243. B) PLEC peptide, 4459-4474. C) PAK1 peptide, residues 88-103.

### List of supplemental tables

**Table S1:** Peptides used for peptide pulldowns.

**Table S2:** Peptide pulldown results with RNF115_69-84_ and NUMB_616-631_ peptides.

**Table S3:** Peptides for PRISMA.

**Table S4:** Identified proteins from peptide pulldown from MS analysis.

**Table S5:** T-test results of identified proteins after peptide pulldown from MS analysis.

**Table S6:** Interaction confidence SAINT-scoring.

**Table S7:** Comparison of enriched proteins for PRISMA using GLUT1 and SOS1 peptides.

